# Proteomic Analysis Reveals Widespread Regulation of Substrate Protein Abundance by O-fucosylation and O-GlcNAcylation

**DOI:** 10.64898/2026.01.16.700008

**Authors:** Sumudu S Karunadasa, Andres V Reyes, TaraBryn S Grismer, Ruben Shrestha, Danbi Byun, Shane Carey, Weimin Ni, Shou-Ling Xu

## Abstract

O-glycosylation of nucleocytosolic proteins by the Arabidopsis enzymes SPINDLY (SPY; O-fucosyltransferase) and SECRET AGENT (SEC; O-GlcNAc transferase) is essential for plant growth and development, yet the scope of their substrates and regulatory impact remains poorly defined. Here, we combined TurboID-based proximity labeling with quantitative proteomics to systematically map the SPY interactome and determine how SPY- and SEC-dependent modifications influence protein abundance. A functional SPY-TD enriched 221 proxiome proteins, including 80 known O-fucosylated substrates and 141 new interactors. The SPY-TD proxiome is enriched in nuclear pore components, chromatin regulators, transcription factors, and RNA-processing proteins. Integration with O-fucose and O-GlcNAc datasets yielded a comprehensive Arabidopsis SPY/SEC (At-S/S) protein list of 886 candidates. We quantified proteome-wide changes in *spy* single mutants and inducible *spy sec* double mutants. Loss of SPY alone caused selective stabilization or destabilization of targets, whereas combined SPY/SEC depletion triggered widespread, synergistic protein abundance changes, particularly affecting nucleoporins, transcriptional regulators, and RNA-binding proteins. Integration with ubiquitination datasets revealed extensive overlap, supporting potential crosstalk between O-fucosylation, O-GlcNAcylation, and ubiquitin-mediated protein turnover. Together, our study establishes proximity labeling as a powerful strategy to define plant O-glycosylation networks and reveals dual, context-dependent roles of SPY and SEC in controlling protein homeostasis and stress-responsive pathways.

## Introduction

In contrast to animals, which rely on a single nucleocytosolic O-GlcNAc transferase (OGT), plants utilize two homologous enzymes: SPINDLY (SPY) and SECRET AGENT (SEC)(Olszewski et al., 2010; Sun, 2021; Mutanwad and Lucyshyn, 2022; Aizezi et al., 2025). While SEC functions as an O-GlcNAc transferase similar to its animal counterparts, SPY has been uniquely characterized as an O-fucose transferase (Zentella et al., 2017; Bi et al., 2023; Zentella et al., 2023). Originally identified in genetic screens for enhanced gibberellic acid (GA) responses, SPY is now known to be a versatile regulator of plant development. It acts as a repressor of GA-mediated processes—such as seed germination, shoot elongation, and flowering—and plays established roles in cytokinin signaling, circadian rhythms, root and style development, immunity, and abiotic stress responses (Tseng et al., 2004; Qin et al., 2011; Steiner et al., 2012; Cui et al., 2014; Steiner et al., 2016; Zhang et al., 2019; Mutanwad et al., 2020; Wang et al., 2020; Jiang et al., 2024).

Conversely, the *sec* single mutant displays a minimal phenotype, suggesting functional redundancy with SPY (Xing et al., 2018; Aizezi et al., 2025). However, this redundancy is vital; the simultaneous loss of both enzymes in the *spy sec* double mutant results in embryonic lethality, underscoring their critical and overlapping functions during early development (Hartweck et al., 2002; Hartweck et al., 2006; Shrestha et al., 2024).

Recent proteomic studies have begun to map the O-glycosylation landscape in plants (Xu et al., 2017; Bi et al., 2023; Li et al., 2023; Wu et al., 2023; Zentella et al., 2023). Using Aleuria aurantia lectin (AAL) and lectin weak affinity chromatography (LWAC), we previously identified 467 O-fucosylation and 489 O-GlcNAc substrates in *Arabidopsis*, with 184 shared substrates. These substrates are central to transcription, RNA processing, and hormone signaling. Despite these advances, current data likely underestimate the true breadth of the plant O-glycosylome, particularly when compared to the thousands of substrates documented in animal systems(Hou et al., 2025). Furthermore, while individual studies show that these modifications influence protein activity, localization, and stability—such as the contrasting effects of O-fucosylation on TCP14 and PRR5 stability—a systematic understanding of how these sugar modifications broadly regulate protein homeostasis remains elusive (Steiner et al., 2012; Zentella et al., 2016; Zentella et al., 2017; Wang et al., 2020; Steiner et al., 2021; Xu et al., 2023a; Jiang et al., 2024; Wang et al., 2025).

To address these gaps, we expanded the SPY interactome using TurboID-based proximity labeling coupled with mass spectrometry (PL-MS). PL-MS is a highly sensitive approach capable of capturing transient interactions that are often missed by traditional affinity purification methods (Mair et al., 2019; Samavarchi-Tehrani et al., 2020; Kim et al., 2023; Xu et al., 2023b; Shrestha et al., 2025). This was particularly advantageous given the low endogenous abundance of SPY and the transient nature of enzyme-substrate interactions. Through optimized enrichment methods, we identified 220 SPY interactors, including 141 previously unreported candidates.

To circumvent the challenge of embryonic lethality and investigate the synergistic roles of these enzymes post-germination, we developed an inducible double mutant system (Shrestha et al., 2025). By introducing a Dexamethasone (Dex)-inducible short hairpin SEC RNAi construct into the *spy-4* background, we generated a line (*shSEC RNAi spy-4*) that allows for the conditional depletion of both glycosyltransferases. While these plants resemble *spy-4* under control conditions, Dex induction (10 µM) triggers a progressive developmental arrest—marked by reduced root growth and leaf discoloration—culminating in seedling lethality by day 7.

Using this inducible system, we performed large-scale quantitative proteomics and parallel RNA-seq profiling across WT, *spy-4*, *sec-5*, and *spy sec* mutants. This integrated approach, quantifying over 10,000 proteins, revealed widespread abundance shifts in *spy-4* that were significantly exacerbated in the *spy sec* double mutant, confirming a deep synergistic interaction. By cross-referencing these findings with public and in-house ubiquitination datasets, we demonstrate that SPY and SEC jointly modulate the stability of numerous substrates, pointing to a sophisticated regulatory crosstalk between O-glycosylation and ubiquitination. Together with the TurboID-derived substrate list, this work provides a foundational, publicly accessible resource for exploring the intersection of sugar modifications and protein homeostasis in plants.

## Results

### Optimized SPY-TD pipeline increases sensitivity and identifies new and known interactors

To expand the identification of SPY interactors, including both substrates and potential regulators, we first performed immunoprecipitation coupled with mass spectrometry (IP-MS) using SPY-GFP (Swain et al., 2002) as bait (Fig. S1A). This analysis identified only four significantly enriched protein groups: SPY itself; two known substrates, transcription factor TCP14 (Steiner et al., 2012) and AT4G22360 (a SWIB complex BAF60b domain-containing protein previously identified via AAL enrichment (Bi et al., 2023); and a novel interactor, AT1G07660, which encodes histone H4. Because many SPY targets are low-abundance nuclear proteins and their interactions are expected to be weak and transient, we sought a more sensitive approach and utilized the TurboID-based proximity labeling method to comprehensively map the SPY interactome (Samavarchi-Tehrani et al., 2020; Xu et al., 2023b).

We generated SPY-TurboID-VENUS (SPY-TD) line, expressed under the *SPY* native promoter in the *spy-3* mutant background (Fig. 1A). To ensure that the SPY-TD fusion protein is fully functional *in planta*, we examined its ability to complement the *spy-3* mutant phenotypes. The *spy-3* mutant exhibits a characteristic smooth leaf phenotype (Fig. 1B) and is resistant to the gibberellin biosynthesis inhibitor paclobutrazol (PAC), allowing it to germinate in its presence (Fig. 1C)(Jacobsen and Olszewski, 1993; Greenboim-Wainberg et al., 2005). As expected, the SPY-TD lines completely rescued the phenotypes, restoring leaf serration and PAC sensitivity to wild-type level (Fig. 1B,C). These results show that the SPY-TD fusion protein maintains full biological activity. For use as a negative control, TurboID fused to YFP (TD-YFP) was expressed under the same native *SPY* promoter (Fig.1A), and a line with expression levels similar to SPY-TD was selected for further analysis. Confocal microscopy confirmed that both SPY-TD and TD-YFP are localized in the nucleus and cytosol (Fig. 1D).

**Figure 1.**
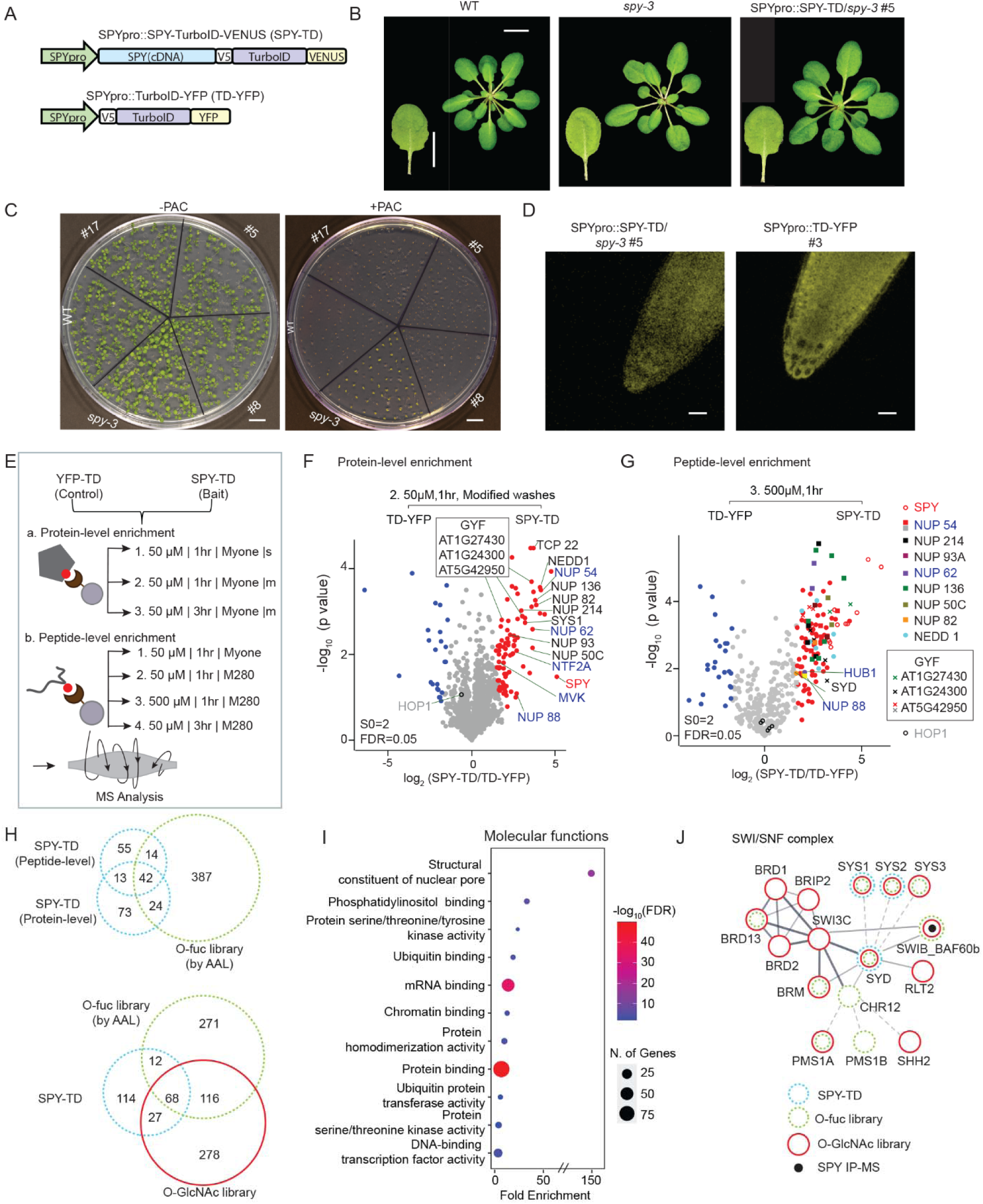
Optimized SPY-TD pipeline increases sensitivity and identifies new and known interactors. (**A**) Schematic representation of the bait (SPY-TD) and control (TD-YFP) constructs, both driven by the native *SPY* promoter. (**B**-**C**) Functional validation of the SPY-TD fusion. The SPY-TD construct fully complements the *spy-3* mutant to wild-type level, as shown by leaf serration (Scale bar= 1 cm) (B) and paclobutrazol (PAC) sensitivity (C). For PAC assays, 3 independent SPY-TD transgenic lines, WT, and *spy-3* were grown on media with or without PAC (15 μg/ml) and germination was assessed after 10 days. Scale bar= 3 cm. (**D**) Confocal microscopy images showing both bait and control proteins localize to the nucleus and cytosol. Scale bar = 25 µm. (**E**) Schematic overview of the dual protein- and peptide-level enrichment workflow designed to maximize coverage of the SPY proxiome (s: standard wash; m: modified wash). (**F**-**G**) Volcano plots showing the enrichment of the SPY-TD interactome at the protein-level (F) and peptide-level (G). The SPY bait, select known O-fucosylation targets (black), and select novel interactors (blue) are highlighted. Multiple nuclear pore complex components and GYF domain proteins were enriched. While protein-level analysis is more sensitive overall, peptide-level enrichment reveals additional interactors, including SYD and HUB1. Proteins that were either non-enriched or preferentially enriched in the TD-YFP control are shown by gray and light blue, respectively. HOP1 is included as a biotinylated control. (**H**) Venn diagram showing the 220 high-confidence proteins comprising the SPY-TD proxiome. The upper panel shows the overlap between protein-level enrichment, peptide-level enrichment, and O-fuc library (identified via AAL enrichment). The lower panel shows the overlap of SPY-TD proxiome with O-fuc and O-GlcNAc (LWAC-enriched) library. (**I**) Gene Ontology (GO) enrichment analysis of molecular functions associated with the high-confidence SPY-TD proxiome. (**J**) Functional interaction network highlighting the connectivity between SWI/SNF chromatin remodeling complexes and SPY/SEC signaling. The network integrates data from multiple independent strategies, including the O-fuc library and O-GlcNAc library, the SPY-TD proxiome, and SPY-GFP IP-MS.

To assess the biotinylation efficiency of the SPY-TD fusion protein, we treated the SPY-TD line with varying concentration of biotin (0-1000 µM) and analyzed the resulting biotinylation patterns via Western blot using α-streptavidin (Fig. S1B,C). Compared to the wild-type control, three specific biotinylated bands were detected in the SPY-TD lines, including a band corresponding to the predicted size of bait fusion itself. However, neither the intensity of these specific bands nor the overall biotinylation signal increased with higher biotin concentrations. This lack of dose-dependence suggests that SPY may interact only weakly or transiently with most of its partners, or that its interactors are of such low abundance that they remain below the detection limit of α-streptavidin in crude extracts.

To overcome these technical limitations and improve detection sensitivity, we optimized SPY-TD enrichment strategies at both the protein- and peptide- levels (Fig. 1E) (Xu et al., 2023b; Karunadasa et al., 2025a; Karunadasa et al., 2025b). Biotinylated species were captured via streptavidin beads. To maximize the recovery of intact biotinylated proteins, we compared different wash conditions (standard vs. modified) and biotin treatment durations (1 h vs. 3 h). We found that the modified wash protocol combined with a 1 h treatment yielded the most robust results. This optimized approach successfully captured several known O-fucosylation targets, including NEDD1, TCP22, GYF proteins (black font), and various nuclear pore components alongside newly identified interactors (blue font) (Fig. 1F, Fig. S2A-C). Interestingly, extending the biotin treatment to 3 h did not improve enrichment efficiency. Instead, the prolonged duration appeared to increase non-specific labeling in both the SPY-TD bait and the TD-YFP control, thereby masking the specific enrichment of the SPY proxiome (Fig. S2C).

To maximize the recovery of biotinylated peptides, we optimized several parameters, including bead type (MyOne vs. M280), biotin concentration (50 µM vs. 500 µM), and treatment duration (1 h vs. 3 h) (Fig. 1G, Fig. S2D-G, Table S1). We found that a 1 h treatment with 500 µM biotin using M280 beads provided the highest sensitivity. This elevated biotin concentration was well tolerated, as excess biotin was effectively removed during peptide preparations before enrichment. This optimized approach not only recovered known substrates but also revealed additional interactors, including several nuclear pore proteins and HISTONE MONO-UBIQUITINATION 1 (HUB1) (blue font).

We integrated data from seven independent datasets and applied stringent filtering criteria to define the SPY interactome (Fig. 1H, Fig. S2H). To ensure high confidence in our identification, biotinylated peptide spectra were manually inspected for the characteristic diagnostic ion of biotinylated lysine (imoKbiotin-NH₃, MW 310.16, C₁₅H₁₉N₂O₂S) (Fig. S3A–C). Protein-level enrichment proved slightly more sensitive, identifying 152 enriched proteins, while peptide-level enrichment identified 124. Among these, 55 were common to both approaches, while 97 and 69 were unique to the protein- and peptide-level enrichment methods, respectively. In total, these analyses yielded a high-confidence set of 221 SPY-TD–enriched proteins, including SPY itself, 80 previously identified O-fucosylated substrates, and 141 newly identified interactors (Fig. 1H) (Table S1). Of these newly identified interactors, 27 overlap with known O-GlcNAcylated proteins, bringing the total number of shared targets between SPY and SEC to 211 (Fig. 1H).

Gene Ontology (GO) analysis of the SPY-TD–enriched proteins revealed significant overrepresentation of molecular functions related to nuclear pore complex components, mRNA and protein binding, transcription factor and chromatin-binding activities, kinase activity, and ubiquitin-related functions (Fig. 1I). While some of these proteins may function as regulators of SPY, many are likely direct SPY substrates, given their overlap with O-fucose (AAL) and O-GlcNAc libraries or their homology to previously characterized substrates (Fig.1H and Table S1). For example, we identified three SWI/SNF chromatin-remodeling components—SYS1, SYS2 and SYD—which are specific subunits of SPLAYED (SPD)-associated SWI/SNF complexes. Their identification reinforces the link between these complexes to nutrient-signaling pathways, consistent with evidence from independent resources (Fig. 1J).

Together, these results demonstrate that TurboID is a robust and sensitive strategy for identifying SPY targets. The SPY-TD dataset substantially expands the catalog of putative SPY targets, highlighting PL-MS as a powerful tool for target discovery that complements traditional enrichment methods. By integrating the SPY-TD, O-fucose (AAL), and O-GlcNAc datasets, we assembled a comprehensive set of 886 proteins, which we collectively refer to as Arabidopsis SPY/SEC (At-S/S) protein list (Table S1).

### MS Strategy to Quantify Proteins Regulated by SPY

SPY has previously been linked to abundance changes in only a few specific target proteins; however, the broader impact on SPY-mediated O-fucosylation on the proteome remains largely unknown. To address this, we analyzed proteomic profiles in the *spy-4* single mutant and a Dex-inducible *ShSEC RNAi spy-4* double mutant (hereafter referred to as *spy sec* double mutants (Shrestha et al., 2025), alongside wild-type (WT) controls. The use of the double mutant allowed us to reveal regulatory effects that might otherwise be masked by the functional redundancy between SPY and SEC. Notably, because the *sec-5* single mutant exhibits an extremely mild, near-WT phenotype and showed negligible proteome-level differences (Fig. S4A; Table S2); it was excluded from the subsequent TMT analysis.

We employed two complementary quantitative proteomic strategies: Stable Isotope Labeling in *Arabidopsis* (SILIA) (Shrestha et al., 2022) and Tandem Mass Tag (TMT) labeling (Ting et al., 2011; Zecha et al., 2019). SILIA enables two sample multiplexing by growing plants for 14 days in media containing either heavy (^15^N) or light (^14^N) nitrogen, followed by mixing of samples and co-analysis (Fig.2A-B, Fig. S4A). Alternatively, TMT16pro allowed for the simultaneous analysis of up to 16 samples (Fig.2C, Fig. S4B). This approach enabled us to examine proteomic changes at an earlier developmental stage (8-day-old seedlings) and provided higher throughput (Fig.2C-D, 3A; Fig.S4F-G).

**Figure 2.**
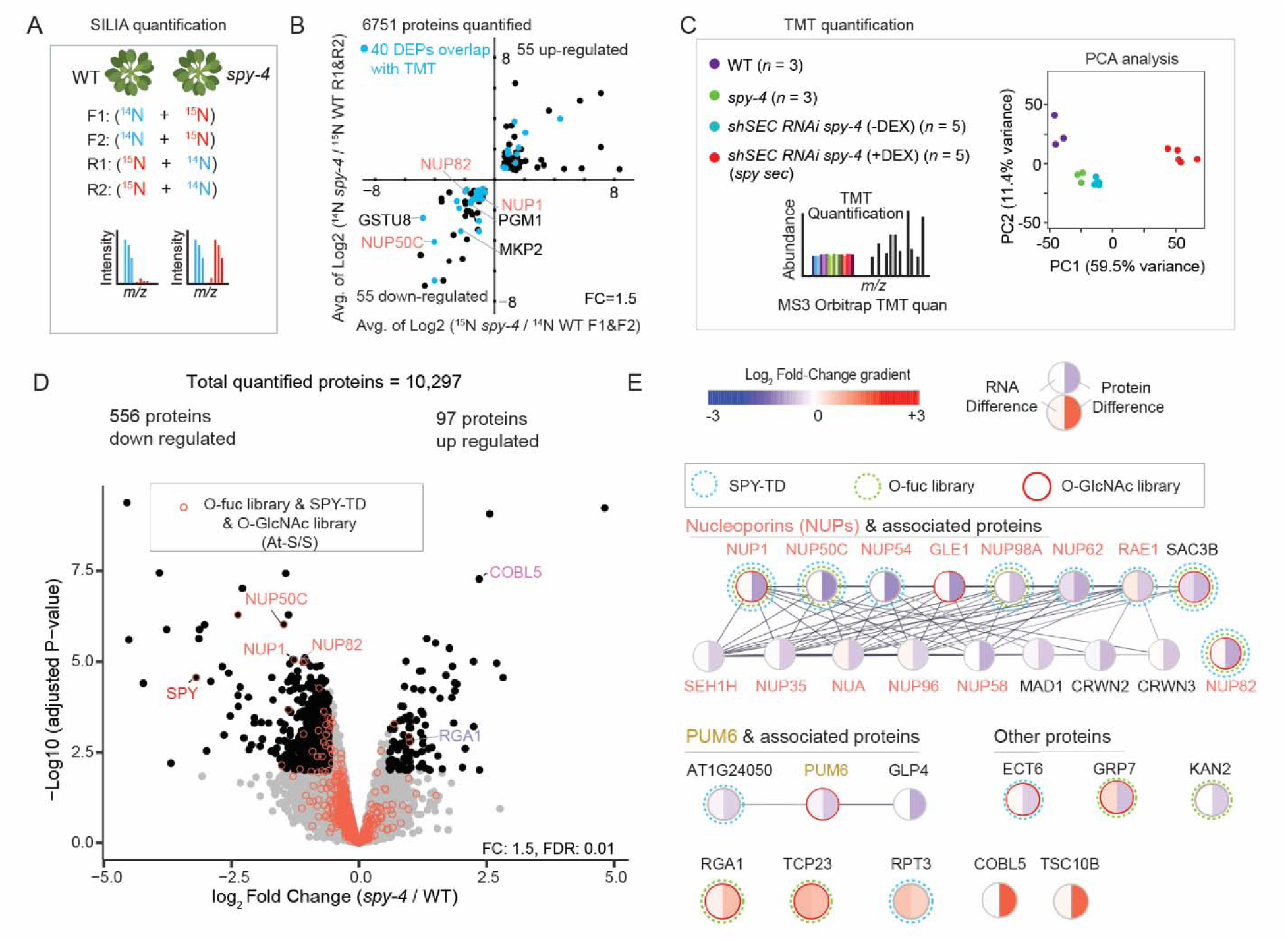
Quantitative proteomic workflow and dual effects of SPY O-fucosylation on protein abundance. (**A**) Schematic of the SILIA quantification comparing 14-day-old WT and *spy-4*. Proteins quantified in at least 3 of 4 biological replicates were retained for downstream analysis. (**B**) Scatter plot showing proteins with altered abundance between WT and *spy-4* as measured by SILIA. The *x*-axis represents the average forward labeling log_2_ ratio (F1&F2), while the *y*-axis represents the average reverse labeling log_2_ ratio (R1&R2). Differentially expressed proteins (DEPs) that were consistently quantified in TMT datasets are highlighted by blue, unchanged proteins are omitted for clarity. Key targets NUP82, NUP1 (NUP136), and NUP50C, are specifically annotated. Additionally, proteins such as GSTU8, PGM1, and MKP2 are highlighted as significantly reduced in the *spy-4* mutant. (**C**) Schematic of TMT quantification and PCA analysis showing high measurement reliability. Distinct clustering is observed for each genotype and treatment group (WT, *spy-4*, *shSEC RNAi spy-4* (± Dex). Notably, mock-treated *shSEC RNAi spy-4* clusters closely with the *spy-4* mutant. (**D**) TMT analysis quantified over 10,000 proteins, identifying 556 down-regulated and 97 up-regulated proteins in the *spy-4* mutant. The volcano plot is overlaid with At-S/S list (red circle). For reference, the SPY bait and targets NUP1, NUP50C, NUP82, and RGA1 are labeled. COBL5 is also highlighted as a protein whose abundance is significantly increased in the *spy-4* mutant. (**E**) SPY-mediated O-fucosylation functions to both stabilize and destabilize substrates, as exemplified by nucleoporins, PUM6, ECT6, and RGA. Protein–protein interactions were visualized using Cytoscape. The left and right halves of each node represent changes in transcript and protein abundance, respectively.

To minimize indirect effects resulting from the severe phenotype of the *spy sec* double mutant, we performed TMT analysis on seedlings grown for five days on standard media followed by three days with Dex-supplemented media (10 µM). This time point corresponds to the initial onset of root growth inhibition in the double mutants. We further optimized the TMT workflow by implementing extensive fractionation to improve depth and data quality. Two mixing strategies were evaluated: equal mixing (Batch I) and unequal mixing (Batch II) (Fig. S4C). The two batches showed strong reproducibility (R² = 0.93, p < 2.2 × 10⁻¹⁶) (Fig. S4D). Batch II, which utilized unequal sample mixing (3x WT relative to mutants) and additional fractionation, enhanced the detection of down-regulated proteins in *spy* and *spy sec* double mutant—including SPY itself—and was therefore selected for the primary analysis.

Using these optimized approaches, we quantified more than 6,700 proteins by SILIA (Fig.2B) and over 10,000 proteins by TMT (Fig.2D, 3A). Principal component analysis (PCA) of the TMT dataset showed tight clustering within genotypes and clear separation between WT, *spy-4*, and *spy sec* double mutant samples (Fig. 2C). Mock-treated *SECi spy-4* samples clustered near *spy-4* but exhibited a slight shift towards the double mutant, indicating mild leaky expression of the RNAi construct, consistent with previous reports (Aoyama and Chua, 1997; Yamaguchi et al., 2015).To determine whether the observed proteomic shifts were driven by the transcriptional regulation, we performed parallel RNA-seq analysis for comparison with the TMT data (Fig. S4B).

### Dual effects of SPY O-fucosylation on protein abundance

To assess the impact of SPY on the Arabidopsis proteome, we first analyzed protein abundance changes in the *spy-4* mutant using SILIA. Applying a 1.5-fold cutoff, we identified 110 differentially expressed proteins (DEPs), consisting of 55 upregulated and 55 downregulated proteins. Notably, 40 of these DEPs were also captured in our TMT-based analysis (Fig. 2B), including three nucleoporins: NUP1, NUP82, and NUP50C. In contrast, no significant abundance difference for these nucleoporins were detected in the *sec-5* mutant using SILIA (Fig. S4A, Table S2).

TMT analysis of WT versus *spy-4* quantified 10,297 proteins, revealing extensive proteomic shifts. We identified 653 DEPs (1.5-fold cutoff), with 556 downregulated and 97 upregulated proteins (Fig 2D, Table S2). Of these, 25 overlap with the At-S/S protein list (red circles). To explore these candidates further, we integrated the RNA-seq and proteomic data using Cytoscape, mapped functional relationships via STRING, and overlaid the At-S/S list to highlight key regulatory nodes (Fig.2E).

We observed a consistent downregulation of nucleoporins and associated proteins at the protein level, while their transcript levels remained largely unchanged. Several RNA-processing factors were also reduced, including the Pumilio protein PUM6 and AT1G24050, an Lsm-domain protein. The YTH domain protein ECT6 also decreased (Xu et al., 2017; Bi et al., 2023; Chen et al., 2023; Shrestha et al., 2024). YTH-domain proteins are critical in multiple steps of RNA metabolism, including splicing, nuclear export, translation, and decay. Additionally, the glycine-rich RNA-binding protein GRP7 and the KANADI transcription factor KAN2 were downregulated. Taken together, these results imply that SPY-mediated O-fucosylation acts to stabilize a specific subset of substrate proteins.

Conversely, the *spy-4* showed increased abundance of several proteins. Unexpectedly, RGA1 levels were elevated. Other known sugar-modified targets, such as TCP23 and the 26S proteasome AAA-ATPase subunit RPT3 also accumulated; however, this was accompanied by increased transcript abundance, complicating the interpretation of these changes. Additional upregulated proteins included the GPI-anchored protein COBL5 and the sphingolipid biosynthetic enzyme TSC10B (3-KDS reductase), although neither is currently known to undergo sugar modification. Collectively, these results demonstrate proteome-wide changes in the *spy* mutant and suggest that SPY-mediated O-fucosylation may both stabilize and destabilize substrate proteins, underscoring its multifaceted role in maintaining protein homeostasis.

### Synergistic effects and widespread proteome changes revealed in inducible *spy sec* double mutants

Due to extensive substrate overlap and functional redundancy between SPY and SEC (Fig.1H) (Aizezi et al., 2025), we reasoned that certain SPY-dependent effects may be masked in single mutants. We therefore analyzed an inducible *spy sec* double mutant. TMT-based analysis identified widespread proteomic shifts, with 3,901 DEPs when compared with WT: 2,765 decreased and 1,136 increased (Fig. 3A, Table S2). Intersecting these with the At-S/S protein list (886 total) showed that 196 of 701 quantified At-S/S proteins were differentially abundant (148 reduced and 48 increased), supporting a direct role for SPY/SEC-dependent modification in regulating protein homeostasis.

**Figure 3.**
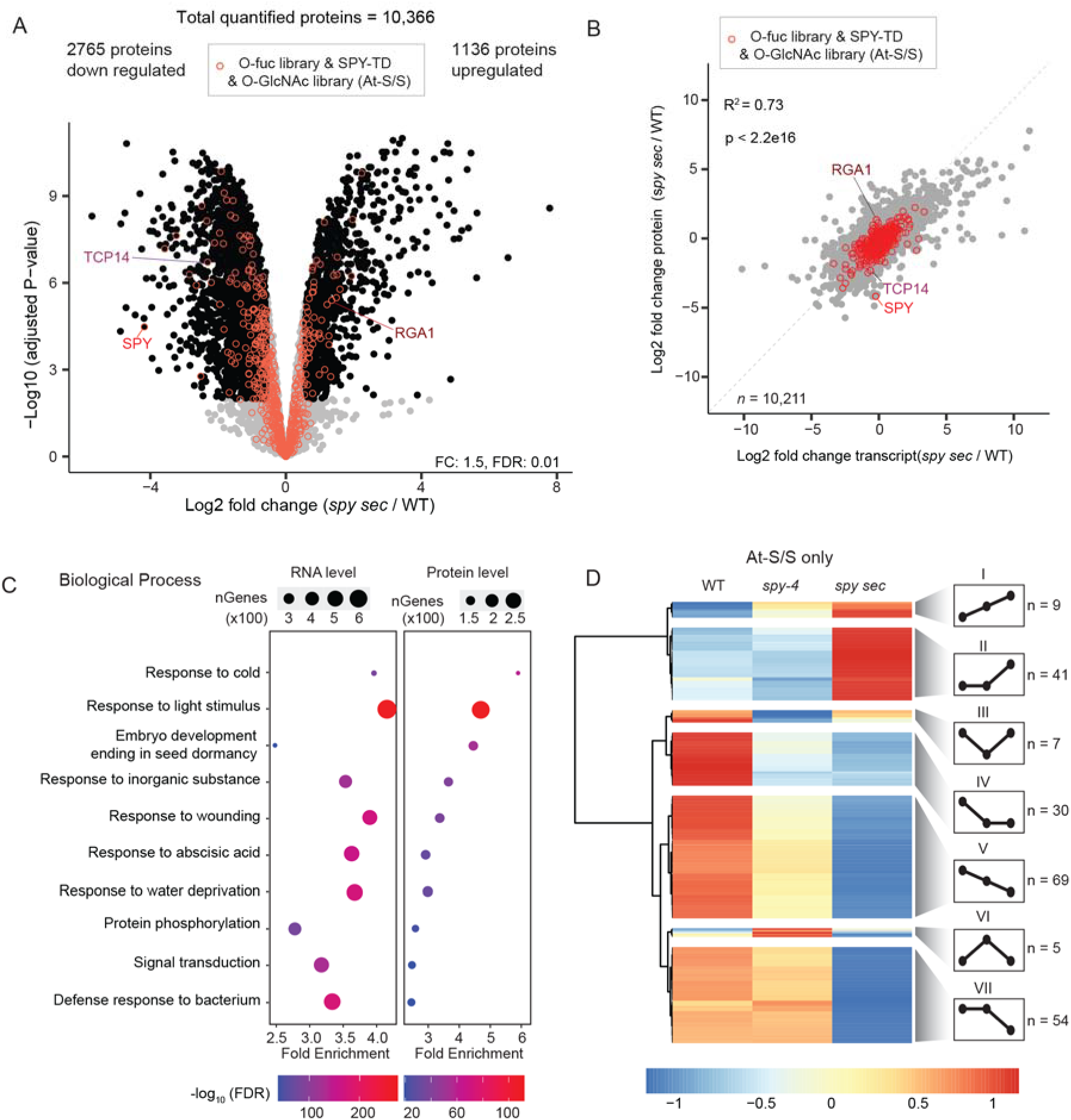
Synergistic effects and widespread proteome changes revealed in inducible *spy sec* double mutants. (**A**) TMT quantification of the *spy sec* double mutant relative to WT. Of over 10,000 quantified proteins, 2,765 were identified as down-regulated and 1,136 as upregulated (FDR=0.01, fold-change = 1.5). Proteins from the At-S/S list are highlighted (red circles), with a substantial fraction (196 of 701 quantified) showing altered abundance (48 up-regulated, 148 down-regulated). SPY, TCP14, and RGA1 are labeled as references. (**B**) Correlation between transcript- and protein-level abundance changes. PCA analysis yielded an R² = 0.73 (p < 2.2 × 10⁻¹⁶), indicating a modest overall correlation. At-S/S proteins are overlaid with red circles. (**C**) GO enrichment analysis of proteins and transcripts showing coordinated regulation. Multiple biological processes are highlighted, including responses to cold, light, wounding, ABA, water deprivation, and bacterial challenge, as well as protein phosphorylation and signal transduction. (**D**) Heatmap of At-S/S proteins showing abundance changes in the *spy-4* single mutant and/or the *spy sec* double mutant. Hierarchical clustering resolved seven expression patterns. Groups I, II, V, and VII (*n* = 173) display synergistic effects, where moderate changes in *spy-4* are markedly intensified in the double mutant. Group IV (*n* = 30) reflects a dominant SPY-dependent effect while Groups III and VI (total *n* = 12) show a restoration effect, consistent with SPY and SEC acting antagonistically on shared targets.

We integrated the RNA-seq data to examine the relationship between transcript and protein abundance changes. Pearson correlation analysis revealed a modest overall correlation (R² = 0.73 (p < 2.2 × 10⁻¹⁶) (Fig.3B). GO analysis identified multiple biological processes with coordinated transcript and protein changes (Fig. 3C), including responses to cold, light stimulus, wounding, ABA, water deprivation, and bacterial challenge. Because O-GlcNAcylation in animals has been implicated in diverse stress-related pathways (Groves et al., 2013; Yang and Qian, 2017; Liu et al., 2021), the broad regulation of stress-associated responses observed in the double mutant suggests a conserved role for the O-glycosylation in stress signaling.

A heatmap comparing proteins from the At-S/S list that showed altered abundance in the *spy* single mutant and/or the *spy sec* double mutants resolved seven distinct expression clusters (Fig. 3D). Group I, II, V, and VII (total *n* = 173) exhibited clear synergistic effects, where proteins showing minimal or modest changes in *spy-4* displayed much more pronounced increases or decreases in the double mutant. In contrast, Group IV (*n* = 30) reflected a dominant SPY-dependent effect, with the double mutant showing shifts similar in magnitude to the *spy-4* single mutant. Groups III and VI (total *n = 12*) revealed a “restoration” effect, where protein levels in the double mutant returned to WT levels, consistent with observations that SPY and SEC can exert opposing effects on shared targets.

### Representative O-glycosylated substrate groups regulated at the protein level

Nucleoporins are among one of the most heavily O-GlcNAcylated proteins in both plants and animals, and many also carry O-fucosylation in plants (Ruba and Yang, 2016; Yoo and Mitchison, 2021; Junod et al., 2025). Several NUPs significantly decreased in *spy-4*, with these reductions becoming more pronounced in the double mutant (Fig.4). Furthermore, seven additional NUPs not altered in *spy-4*—including NUP35, NUP93B, and NUP155—displayed altered abundance in the double mutant. These are not in the current At-S/S list; their coordinated reduction may reflect the co-degradation of compromised protein complexes (Ryan et al., 2017).

**Figure 4.**
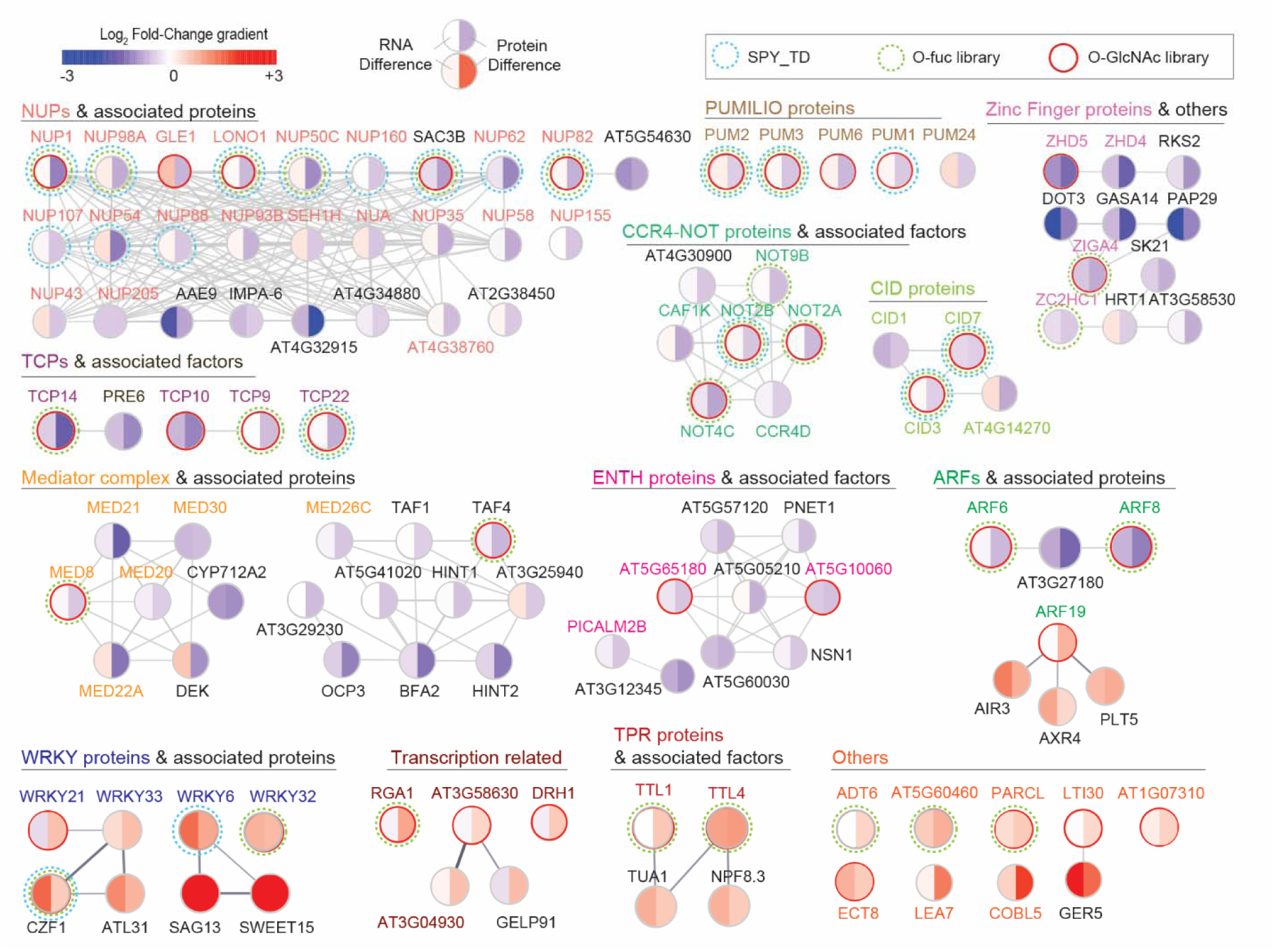
Representative O-glycosylated substrate groups regulated at the protein level. A broad spectrum of O-glycosylated proteins exhibits either stabilization or destabilization in the *spy sec* double mutant. Representative down-regulated groups include nucleoporins, TCP transcription factors, PUMILIO proteins, CCR4–NOT complex components, CID proteins, zinc-finger transcription factors, Mediator complex subunits, and ENTH-domain proteins. In contrast, Auxin Response Factors (ARFs) display bidirectional changes in abundance. Representative up-regulated groups, including WRKY transcription factors, RGA1 and other transcription related proteins, TPR containing proteins, are also shown. For each visualized node, the left half represents transcriptomic changes, while the right half denotes proteomic changes, with nodes overlaid with the At-S/S dataset for reference. The network was generated in Cytoscape using STRING database interactions.

The plant-specific TCP transcription factor family, a known target of sugar modifications, also exhibited heightened sensitivity. TCP14 remained unchanged in *spy-4* but was sharply reduced in the double mutant, alongside TCP9, TCP10, and TCP22. Broader regulatory networks in transcription and mRNA metabolism were similarly affected. Multiple sugar-modified proteins—including PUMILIO RNA-binding proteins, components of the CCR4–NOT complex, CID-interacting proteins (orthologs of human ATXN2), zinc-finger transcription factors, and Mediator subunits—showed significant reductions in the double mutant (Fig. 4). ENTH-domain proteins, which participate in membrane dynamics (Feng et al., 2022) and known O-GlcNAcylation targets, were also consistently reduced. Hormone-related regulators were affected as well: ARF6 and ARF8 decreased, whereas ARF19 increased, underscoring the diverse and sometimes opposing effects of altered sugar modifications on protein abundance.

In contrast, a distinct set of proteins accumulated in the double mutant. WRKY family members and associated regulators increased at both the transcript and protein levels. RGA1, already elevated in *spy-4*, rose further in the double mutant. Other stabilized factors included transcriptional regulators (e.g., AT3G58300), the DEAD-box RNA helicase DRH1, and TTL1. Intriguingly, several intrinsically disordered proteins (IDPs)—including LTI30 and AT1G07310—accumulated, which is notable given that sugar modifications often occur within intrinsically disordered regions. Arogenate dehydratase 6 (ADT6), a plastid-localized enzyme and known O-fucosylated substrate, also showed increased protein levels.

### Potential Crosstalk of O-GlcNAcylation and O-fucosylation with Ubiquitination

In animals, O-GlcNAcylation modulates protein ubiquitination, either enhancing or inhibiting it depending on the substrate (Yang et al., 2006; Hardiville et al., 2010; Park et al., 2010; Srikanth et al., 2010; Ruan et al., 2013). To investigate similar crosstalk in plants, we assessed whether proteins with altered abundance in the *spy sec* double mutant are also ubiquitinated.

We generated an in-house ubiquitination dataset and performed extensive meta-analysis of ten public datasets (Maor et al., 2007; Saracco et al., 2009; Book et al., 2010; Kim et al., 2013; Walton et al., 2016; Aguilar-Hernandez et al., 2017; Grubb et al., 2021; Ma et al., 2021; Berger et al., 2022; Song et al., 2024), identifying a combined total of 7,448 ubiquitinated proteins (Fig. 5A, Fig.S5A, Table S3). Notably, many downregulated proteins in the *spy sec* mutant, including NUPs, TCPs and kinases like ERECTA and PGK1, are ubiquitinated (Fig.5A, Fig.S5B-C). Conversely, upregulated sugar-modified proteins, such as RGA1, also overlap with the ubiquitinated proteome (Fig.5A, Fig.S6A-B). Given that over 5,000 sugar-modified targets exist in animals, we anticipate that many plant substrates remain unidentified but likely exhibit altered abundance in our mutants. GO enrichment analysis of ubiquitinated proteins with altered abundance revealed enrichment in pathways such as chloroplast protein import and chlorophyll biosynthesis, likely underlying the leaf discoloration phenotype of the double mutant (Fig.5A). We also observed enrichment in proteins involved in embryo development and seed dormancy, potentially explaining the lethality of the *spy sec* mutant. In summary, our data suggest that SPY and SEC synergistically regulate substrate abundance. The overlap with the ubiquitinated proteome supports a model of crosstalk between O-GlcNAcylation, O-fucosylation, and ubiquitination (Fig. 5B). Furthermore, given the association of SPY with histones and chromatin-binding proteins (Fig. 1J), we hypothesize that SPY and SEC may also directly regulate transcription (Fig.5B).

**Figure 5.**
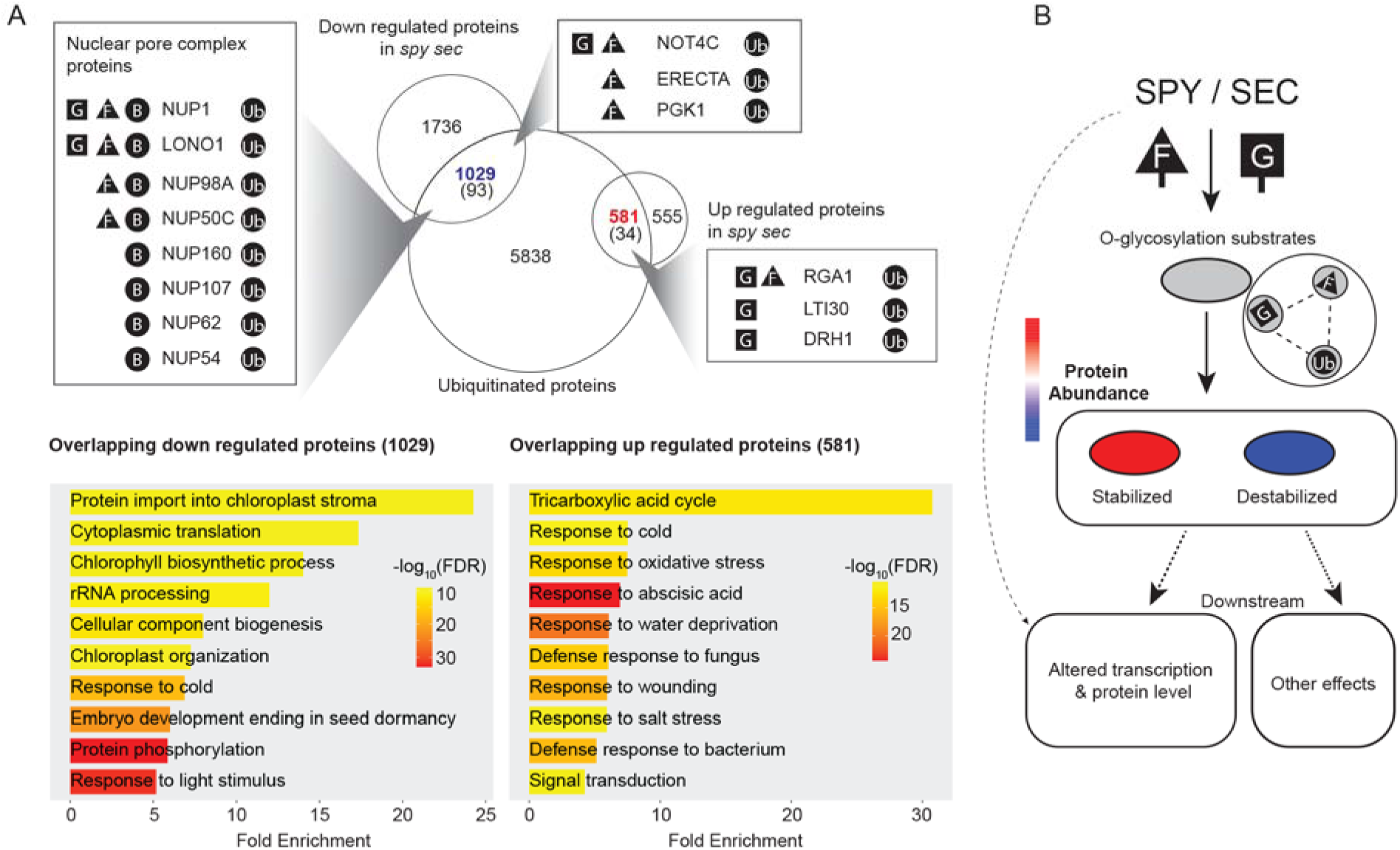
Potential crosstalk between O-GlcNAcylation, O-fucosylation, and ubiquitination, and a proposed model for SPY/SEC-mediated regulation. (**A**) Overlap between differentially expressed proteins (up-regulated and down-regulated) in the *spy sec* double mutant and a curated list of known ubiquitinated proteins. Values in brackets indicate the number of overlapping proteins present in the At-S/S list, with representative functional subsets highlighted in inset boxes. Lower panels show Gene Ontology (GO) biological process enrichment for the overlapping protein sets. (**B**) Proposed model for SPY- and SEC-mediated regulation of protein transcription and abundance. SPY and SEC modulate substrate stability through O-fucosylation and O-GlcNAcylation, potentially via biochemical crosstalk with the ubiquitin-proteasome system. These modifications lead to coordinated changes in transcript levels and protein accumulation, ultimately driving the pleiotropic phenotypes observed in *spy* and *spy sec* mutants. Furthermore, given the association of SPY with various chromatin-remodeling proteins, SPY (and SEC) may also directly regulate gene transcription.

## Discussion

Using PL-MS, we identified 221 SPY-TD enriched proteins, thereby considerably expanding the known SPY substrate repertoire beyond the 80 proteins shared with existing AAL datasets (Fig.1H). The combined protein- and peptide-level enrichment approaches yielded complementary information and defined a high-confidence SPY-associated proteome, which is likely to comprise both direct substrates and proteins that interact with or regulate SPY activity. While SEC-TD was also investigated, its analysis was hindered by low bait expression relative to the TD-YFP control under the same native SEC promoter (data not shown). This unfavorable expression ratio resulted in excessive background labeling that obscured potential substrates; consequently, we focused our downstream analysis on the robust and high-quality SPY-TD dataset.

Through the integration of quantitative proteomics (SILIA/TMT) and transcriptomics, we mapped the proteomic landscape regulated by SPY and SEC. We found that while SPY possesses unique autonomous targets, the two enzymes predominantly operate in synergy to maintain proteome homeostasis (Fig. 3D). Most notably, we observed extensive shifts in protein abundance that did not correlate with changes in transcript levels; the quantification establishes O-fucosylation and O-GlcNAcylation as key post-translational modifiers of protein stability. These data support a model where nucleocytoplasmic glycosylation acts as a sophisticated rheostat, controlling protein turnover across the proteome. Importantly, our data provide the first large-scale evidence that SPY and SEC broadly govern protein abundance in plants. This mechanism appears to be evolutionarily conserved, as SPY has recently been shown to similarly regulate protein homeostasis in *Toxoplasma gondii* (Tiwari et al., 2025).

A striking feature of our dataset is the enrichment of nucleoporins among proteins regulated by both SPY and SEC. While O-GlcNAcylation is a well-established modulator of nuclear pore activity and stability in metazoans (Ruba and Yang, 2016; Yoo and Mitchison, 2021; Junod et al., 2025), our findings extend this paradigm to the plant kingdom. Specifically, we demonstrate that SPY and SEC co-regulate nucleoporin abundance, with SPY emerging as the dominant regulator in this biological context. In addition, we found that numerous proteins involved in transcriptional regulation—including members of the TCP and PUMILIO families, the CCR4–NOT complex, and Mediator subunits—are subject to O-glycosylation–dependent control at the protein level. Notably, although previous studies reported that TCP14 stability is SPY-dependent under overexpression conditions (Steiner et al., 2012; Steiner et al., 2016; Steiner et al., 2021), our data show that endogenous TCP14 abundance is significantly altered only in the *spy sec* double mutant. This observation suggests a compensatory buffering mechanism at physiological expression levels, in which loss of one glycosyltransferase is partially offset by the other, thereby maintaining homeostatic control over key transcription factors. We observed complex and divergent changes in ARF abundance in the *spy sec* double mutant, characterized by decreased levels of ARF6/8 and increased accumulation of ARF19. Together with the elevated abundance of substrates such as RGA1, these findings demonstrate that altered O-glycosylation can exert opposing effects on protein stability.

Furthermore, the pronounced changes in protein abundance—largely uncoupled from transcript levels—suggest a functional interplay between O-glycosylation and the ubiquitin–proteasome system. In metazoans, O-GlcNAcylation modulates protein ubiquitination in a substrate-dependent manner, either promoting or inhibiting protein turnover (Yang et al., 2006; Hardiville et al., 2010; Park et al., 2010; Srikanth et al., 2010; Ruan et al., 2013). Consistent with this paradigm, a substantial fraction of proteins altered in the *spy* or *spy sec* mutants are known ubiquitination targets, suggesting that O-glycosylation may directly influence protein half-life in plants.

While our inducible *shSEC* RNAi *spy-4* line displays a more severe phenotype than previously reported double mutants (e.g., *a2* SEC RNAi *spy-3*) (Jiang et al., 2024; Shrestha et al., 2025), SEC may not be fully depleted within three days of Dex treatment. Consequently, our analysis likely represents a conservative estimate of the SPY/SEC-regulated proteome. Interestingly, our data suggest that point toward a previously unrecognized homeostatic feedback loop between these two enzymes, as SEC knockdown led to a compensatory increase in SPY transcript levels (Supplementary Fig. S4E).

Beyond regulating protein stability, the broad enrichment of stress-related pathways in the double mutant (Fig. 3C) suggests a conserved role for O-glycosylation in plant stress signaling, echoing its role in metazoans (Groves et al., 2013; Yang and Qian, 2017; Liu et al., 2021). Our identification of SPY associations with histones and chromatin-remodeling factors—via both IP-MS and PL-MS—further suggests that SPY and SEC may directly modulate the transcription of many genes. This is highly analogous to mammalian OGT, which is enriched at key regulatory chromatin regions (Gambetta et al., 2009; Dey et al., 2012; Chen et al., 2013; Vella et al., 2013) and forms stable complexes with transcriptional regulators such as HCF1 and TET enzymes (Deplus et al., 2013; Vella et al., 2013; Hrit et al., 2018). Determining whether SPY and SEC exhibit similar genome-wide co-occupancy to regulate gene expression in plants remains a critical frontier for future investigation.

## Materials and Methods

### Construct generation and transformation

The pENTR1A-SPY construct was generated by amplifying the full-length coding sequence of SPY using PCR, which was then cloned into the pENTR1A vector (Thermo Fisher Scientific). Approximately 2 kb upstream of the SPY ATG start codon, which encompasses the promoter and 5’-untranslated leader (Swain et al., 2001) was amplified and cloned into the pENTR5 vector gateway clone to generate the SPY promoter entry clone (pENTR5-SPYpro). Three entry vectors (pENTR5-SPYpro, pENTR1A-SPY, pDONR_P2R-P3_R2-Turbo-mVenus-STOP-L3) were recombined with destination vector pB7m34GW (Mair et al., 2019) to generate the final expression vector SPYpro::SPY-TD-mVENUS (abbreviated as SPY-TD). The control TurboID-YFP entry vector (pENTR_L1-Turbo-YFP-STOP-L2) was made by applying site-directed mutagenesis to introduce a stop codon right before the NLS of the entry clone (pENTR_L1-Turbo-YFP-NLS-STOP-L2) (Mair et al., 2019) to eliminate the NLS signal sequence. The control construct SPYpro::TD-YFP (abbreviated as TD-YFP) was generating by recombining two entry vectors (pENTR5-SPYpro, pENTR_L1-Turbo-YFP-STOP-L2) with destination vector R4pGWB501 to generate the final expression vector. SPY-TD was transformed into the *spy-3* mutant (point-mutation allele) and TD-YFP transformed into wild type via the floral-dip method (Clough and Bent, 1998) and selected based on BASTA and hygromycin, respectively. Complemented SPY-TD lines and TD-YFP lines with a similar expression level to that of SPY-TD were chosen for further analysis. *ShSECRNAi* construct and related line was described in (Shrestha et al., 2025).

### Complementation phenotype examination

For PAC resistance test, WT, *spy* mutant, and three independent SPY-TD transgenic lines were germinated in the presence of the gibberellin biosynthesis inhibitor, paclobutrazol (PAC). Sterilized seeds were plated on ½ Murashige & Skoog (MS) media (2.17g/L MS, 6 g/L Phytoblend, pH 5.8) (control) and ½ MS medium supplemented with 15 μg/ml PAC (treatment). The seeds were placed in 4 °C cold room for 2 days for stratification then transferred to a chamber at 22 °C with a 24h-light (light intensity of 82 μmol m^−2^ s^−1^) for 10 days. Germination in the presence of PAC was assessed in the transgenic lines compared to wild type and *spy-3* mutant. For leaf serration analysis, phenotypic analysis of SPY-TD transgenic lines was performed based on the presence or absence of leaf serration. Leaves were assessed from three weeks old plants grown in greenhouse conditions with 100 µmol m^−2^ s^−1^ light intensity.

### Confocal microscopy

Confocal microscopy images of Arabidopsis seedlings expressing TurboID constructs were taken with a Leica SP8 microscope. Images were processed using Adobe photoshop software.

### Streptavidin immunoblot

10- or 5-day-old Arabidopsis seedlings were treated with 0–1000 µM biotin for 1 h. 40 mg of tissue per sample were flash-frozen, and proteins were extracted in 2× SDS loading buffer (1:3 w/v). Proteins were separated by SDS-PAGE, transferred to 0.2 µm PVDF membranes (BioRad), blocked with 3% BSA, and probed with anti-streptavidin antibody (1:2000, Thermo Fisher Scientific).

### Plant growth and biotin treatment for TurboID experiments

SPY-TD and TD-YFP seedlings were grown on ½ MS medium, cold-treated for 3 days, then incubated vertically under 24 h light at 22 °C for 10 days. Ten-day-old seedlings (1–2 g, three replicates per condition) were incubated in 50–500 µM biotin for 1 h at room temperature, washed, patted dry, flash-frozen in liquid nitrogen, ground into a fine powder, and stored at –80 °C until protein extraction.

### Immunoprecipitation of the SPY Interactomes (IP-MS)

The SPY-GFP (Swain et al., 2002) and TAP-GFP seedlings (Shen et al., 2008) were grown for 7 days at 21°C under constant light on ½ MS medium. Tissues were harvested, flash-frozen in liquid nitrogen, ground into a fine powder, and stored at −80°C. Immunoprecipitation was performed as previously described with slight modifications (Ni et al., 2013). Briefly, proteins were extracted in MOPS buffer (100 mmol/L MOPS, pH 7.6, 150 mmol/L NaCl, 1% (v/v) Triton X-100, 1 mmol/L PMSF, 2× Complete protease inhibitor cocktail, and PhosStop cocktail (Roche)), centrifuged, and filtered through two layers of Miracloth, followed by incubation with a modified version of LaG16-LaG2 anti-GFP nanobody conjugated to Dynabeads (Invitrogen) for 3 h at 4°C. Incubation was followed by four 2-minute washes with immunoprecipitation buffer and eluted with 2% (w/v) SDS buffer containing 10 mmol/L tris(2-carboxyethyl) phosphine (TCEP) and 40 mmol/L chloroacetamide at 95°C for 5 minutes. Eluted proteins were separated by SDS-PAGE. After Colloidal Blue staining, the whole lane of protein samples was excised in three segments and subjected to in-gel digestion with trypsin. Three biological replicates were performed.

### Biotinylated species enrichment at protein-level and peptide-level

Biotinylated species were enriched at the protein or peptide level as described (Karunadasa et al., 2025a; 2025b). For protein-level enrichment, 200 µL MyOne Streptavidin C1 Dynabeads (Invitrogen) were used following established protocols (Branon et al., 2018; Mair et al., 2019), with variables including standard versus modified washes (Karunadasa et al., 2025b) and biotin incubation times of 1 h or 3 h. For peptide-level enrichment, variables included bead type (MyOne T1, #65602, or M-280, #11206D, Invitrogen), biotin concentration (50 µM vs 500 µM), and biotin treatment duration at 50 µM (1 h vs 3 h).

### SILIA sample preparation

The SILIA proteome quantification of WT relative to *spy-4* plants and WT relative to *sec-5* plants were performed as described in.(Shrestha et al., 2022). Briefly, WT and mutant lines (*spy-4, sec-5)* were grown for 14 days on Hoagland’s medium containing ^14^N or ^15^N (1.34 g/L Hoagland’s No. 2 salt mixture without nitrogen, 6 g/L Phytoblend, and 1 g/L KNO_3_ or 1 g/L K^15^NO_3_ (Cambridge Isotope Laboratories), pH 5.8, 1% sucrose). Plates were placed vertically in a growth chamber under constant light conditions at 21–22 °C. Whole plant tissues were harvested in liquid nitrogen. Proteins were extracted from eight samples individually using 2X SDS sample buffer (plant tissue mass (mg): Buffer (μL) ratio = 1:3) and mixed as follows: two forward samples F1 and F2 (^14^N WT/^15^N mutant,) and two reverse samples R1 and R2 (^14^N mutant/^15^N WT). Proteins were separated by SDS-PAGE, and five gel segments were excised for in-gel trypsin digestion. The resulting peptide mixtures were desalted using C18 ZipTips (Millipore) and analyzed by liquid chromatography–mass spectrometry (LC-MS).

### TMT sample preparation

For tandem mass tag (TMT)–based quantitative proteomic analysis, Arabidopsis thaliana wild-type (WT), *spy-4*, and *shSEC RNAi spy-4* seedlings were used. WT and *spy-4* seedlings were grown vertically on half-strength Murashige and Skoog (½ MS) medium for 8 days before harvesting. *shSEC RNAi spy-4* seedlings were grown on ½ MS medium for 5 days and then transferred to plates supplemented with either DMSO (mock) or 10 µM dexamethasone (Dex) for 3 additional days before harvesting. All seedlings were grown under constant light conditions.

For proteomic analysis, 50 mg of tissue from three biological replicates of WT, three biological replicates of *spy-4*, and five biological replicates each of mock- and Dex-treated *shSEC RNAi spy-4* seedlings were used. Protein extraction was performed as described as (Xu et al., 2017), with a slight modification to the extraction buffer composition (100 mM Tris-HCl, pH 8.0; 2% [w/v] SDS; 5 mM EGTA; 10 mM EDTA; 1 mM PMSF; and 2× protease inhibitor cocktail, Roche). Extracts were reduced with TCEP, alkylated, and digested with trypsin, followed by peptide desalting using Sep-Pak C18 cartridges (Waters). TMT labeling was performed using a TMT 16-plex kit (Thermo Fisher Scientific, Cat# A44520).

TMT 16-plex reagents were reconstituted in anhydrous acetonitrile (0.01 mg/μL). Approximately 200 μg of peptides from each sample were dissolved in 60 μL of 50 mM HEPES buffer and labeled with 40 μL of TMT reagents. Reactions proceeded for 1 h at 25 °C with shaking at 700 rpm and were quenched with 5 μL of 1 M Tris (pH 8). Peptides were then acidified with 45 μL of 10% FA in 10% acetonitrile. The final reaction contained TMT at 4 μg/μL, peptides at 2 μg/μL, and 40% acetonitrile (v/v) (Zecha et al., 2019).

High-pH reversed-phase HPLC fractionation was performed in two batches as follows. In Batch I, all 16 labeled samples were mixed in equal proportions and fractionated, pooled into 12 runs. In Batch II, the proportion of WT samples increased threefold prior to channel pooling, followed by fractionation into 48 runs. High pH reverse-phase chromatography was performed on the Vanquish Flex system (ThermoFisher) equipped with a 4.6×150-mm Gemini 5μm C18 column for TMT-labeled peptides. Peptides were loaded onto the column in 100 μL of buffer A (20 mM ammonium formate, pH 10). Buffer B consisted of buffer A with 90% (vol/vol) acetonitrile. Peptides were separated at 0.5 mL/min using high-pH reversed-phase gradients. Common steps included column equilibration at 1% B and sequential increases to 50–70% B, 70–100% B, and 100% B hold. Fractions were collected every 1 min. Collected fractions were vacuum-dried and subjected to LC/MS/MS analysis.

### diGly proteomics analysis to identify ubiquitinated proteins *in vivo*

Metabolically labeled Arabidopsis plants with ^14^N and ^15^N isotope were harvested, followed by protein extraction and trypsin digestion as previously described (Xu et al., 2017; Shrestha et al., 2022). Ubiquitinated peptides were enriched using the PTMScan Ubiquitin Remnant Motif (K-ε-GG) Kit (Cell Signaling Technology) according to the manufacturer’s instructions.

### LC-MS/MS analysis

Peptides were analyzed by liquid chromatography–tandem mass spectrometry (LC-MS) on an EasyLC1200 system (Thermo) connected to a high-performance quadrupole Orbitrap Q Exactive (IP-MS samples and SILIA samples) or an Orbitrap mass spectrometer Eclipse (all other samples)(Thermo). Peptides were first trapped using trapping column Acclaim PepMap 100 (75 µm x 2cm, nanoViper 2Pk, C18, 3 µm, 100A), then separated using analytical column AUR2-25075C18A, 25CM Aurora Series Emitter Column (25cm x75 µm, C18, 1.6um) (IonOpticks). The flow rate was 300 nL/min. For peptide-level enrichment samples, peptides were eluted by a gradient from 3 to 10% solvent B (80% (v/v) acetonitrile/0.1% (v/v) formic acid) over 1 min, 10 to 35% solvent B over 105 min, from 35 to 44% solvent B over 15 min, followed by a short wash for 15 min by 90% solvent B. For protein-level enriched samples, peptides were eluted by a gradient from 3 to 5% solvent B over 1 min, 5 to 28% solvent B over 105 min, from 28 to 44% solvent B over 15 min, followed by a short wash (15 min) at 90% solvent B.

For data acquisition on Eclipse, precursor scan was from mass-to-charge ratio (m/z) 375 to 1600 (resolution 120,000; AGC 200,000, maximum injection time 50ms, Normalized AGC target 50%, RF lens(%) 30) and the most intense multiply charged precursors were selected for fragmentation (resolution 15,000, AGC 5E4, maximum injection time 22ms, isolation window 1.4 m/z, normalized AGC target 100%, include charge state=2-8, cycle time 3 s). Peptides were fragmented with higher-energy collision dissociation (HCD) with normalized collision energy (NCE) 27. Dynamic exclusion was enabled for 30s.

For Orbitrap Q Exactive (Thermo), peptides were separated using ES803 (50cm) column. The LC gradient is the same as above. Precursor scan was from mass-to-charge ratio (m/z) 375 to 1600 (resolution 120,000; AGC 3.0E6, maximum injection time 100 ms) and top 20 most intense multiply charged precursors were selected for fragmentation (resolution 15,000, AGC 5E4, maximum injection time 60ms, isolation window 1.0 m/z, minimum AGC target 1.2e3, intensity threshold 2.0 e4, include charge state =2-8). Peptides were fragmented with higher-energy collision dissociation (HCD) with normalized collision energy (NCE) 27. Dynamic exclusion was enabled for 24s.

### SILIA data Search and quantification/filtering

MS/MS data were searched against the TAIR database using Protein Prospector (Shrestha et al., 2022) with a precursor mass tolerance of 10 ppm and fragment tolerance of 20 ppm. Carbamidomethyl (C) was set as a fixed modification, and variable modifications included protein N-terminal acetylation, peptide N-terminal Gln conversion to pyroglutamate conversion, and Met oxidation. SILIA comparisons (WT vs. *spy-4* and WT vs. *sec-5*) included two forward (F1, F2) and two reverse (R1, R2) replicates. ^14^N /^15^N ratios were transformed to represent mutant/WT and normalized to the mean of the top 100 proteins in each replicate. The quantitative ratios were merged into one table by protein accession and then log₂-transformed. To reduce contamination and noise, data were filtered by: (1) retaining proteins quantified in ≥3 replicates, and (2) selecting proteins with log₂-fold changes [FC]≥1.5 (or consistently within –1.5 to 1.5 for non-regulated proteins). Ubiquitinated data were searched using Protein Prospector, with di-Gly on lysine specified as variable modification.

### Data search and label free quantification (LFQ)

Raw data were processed using MSFragger (v20.0) for identification against the TAIR10 database (35,386 entries) and LFQ. Search parameters included 10 ppm precursor and 20 ppm MS/MS tolerance, with variable modifications as described above.

For SPY-TD peptide-level enrichment, cleavage was set to trypsin (two missed cleavages) with biotinylation at uncleaved lysines (Biotin(K)) included as a variable modification (max 3 per peptide). For SPY-TD protein-level enrichment, one missed cleavage and two modifications were allowed, excluding the biotin modification; a minimum of two values was required for quantification. LFQ was performed with a 1% FDR at both the peptide and protein levels, enabling the match between runs. For SPY-GFP IP-MS enrichment, data were searched in Maxquant with similar settings.

### Statistics processing using Perseus for LFQ quantification

For protein-level enrichment MSFragger’s ‘combined_proteins’ was input to Perseus, also with LFQ intensities specified as ‘Main’ columns. Decoys and/or contaminants were removed, then the LFQ intensities were log2 transformed and quality checked with histograms. Non-reproducible identifications were removed (keep only rows with at least 3 valid values in either SPY-TD replicates or TD-YFP replicates), then imputation from a normal distribution was performed to replace remaining missing values. A two-sided T-test with S0=0.5, and FDR=0.05 or 0.1 were used for protein level (See filtering steps).

For peptide-level enrichment MSFraggers’s ‘combined_modified_peptides’ was loaded into Perseus v2.0.10, with LFQ intensities specified as ‘Main’ columns. The dataset was filtered to retain only biotinylated peptides for further statistical analysis. Statistical significance and enrichment for each identified peptide were determined using a two-sided Student’s t-test (S0=2, FDR=0.05). To facilitate protein-level reporting, biotinylated peptides were collapsed by protein loci, and the mean LFQ intensity was calculated. A second two-sided t-test was then performed on these mean values to assess protein-level significance. Furthermore, the median difference of all quantified biotinylated peptides was reported to characterize the enrichment distribution of the peptides associated with each protein.

### Filtering steps for SPY-TD

Protein and peptide quantification was performed in Perseus using MSFragger outputs. Significant enrichment thresholds were set at S0=2, FDR=0.05 (peptide) and S0=0.5, FDR=0.1 (protein). This identified 221 high-confidence interactors (including SPY) across four categories:

- Category 1 (55 proteins): Enriched at both protein and peptide levels.
- Category 2 (24 proteins): Enriched in ≥ 2 independent protein-level datasets.
- Category 3 (73 proteins): Enriched in one protein-level dataset (fold enrichment ≥ 2, FDR is reduced to 0.05).
- Category 4 (69 proteins): Enriched exclusively at the peptide level.

Category 4 biotinylation was validated via MSviewer, targeting the m/z 310.16 signature ion (ammonia-loss immonium ion of biotinylated Lys) and proline signature ions. Quantification was refined in Skyline (5 ppm MS1 mass accuracy), where peptides were manually curated to ensure high-quality spectra and clear enrichment over YFP controls.

### TMT data analysis

TMTpro raw files were processed using MaxQuant (v2.4.2.0) in MS3 ion reporter mode. Reporter ion intensities were corrected using manufacturer-provided factors without additional normalization. Search parameters included a first-round peptide mass tolerance of 20 ppm, a main search tolerance of 4.6 ppm, and an MS2 match tolerance of 20 ppm. The reporter mass tolerance and isobaric weight exponent were set to 0.003 and 0.75, respectively. Sequences were searched against Arabidopsis thaliana TAIR10 database. For protein-level analysis, proteinGroups.txt and evidence.txt files were analyzed using the MSstatsTMT R package (Huang et al., 2020). Protein summarization was performed via the msstats method with global normalization enabled, while reference normalization and remove normalization channel were disabled. Protein-level quantifications were analyzed across all samples. For proteins detected in all five replicates of one condition but absent in the other, missing values were imputed from a normal distribution (width: 0.3, downshift: 1.8). Differential abundance was assessed using the MSstatsTMT’s groupComparisonTMT function. Proteins with missing p-values were filtered out. If an ambiguous protein group existed with different gene loci, they were discarded. If multiple protein groups existed for a single gene locus, the protein group with the most peptide numbers quantified was kept. Protein groups with an FDR=0.01 and a [FC] ≥1.5 were considered as differentially abundant proteins.

### Network analysis, visualization and heatmaps

Proteins with significantly altered abundance in the *spy-4* and *spy sec* double mutant were visualized using Cytoscape (v3.10.2) (Otasek et al., 2019). Protein–protein interaction data were retrieved via the STRING database plug-in for Arabidopsis thaliana. Transcript- and protein-level fold changes were mapped as node attributes and displayed side by side to facilitate direct comparison. Node colors were scaled by fold-change magnitude and direction. Final network layouts were generated using standard Cytoscape algorithms to visualize functional relationships. Heat maps were generated in R using log₂ fold-change values from the TMT proteomics and RNA-seq datasets relative to wild type. For specified visualizations, hierarchical clustering was applied to identify patterns of co-regulation.

### Ubiquitome compilation and Gene Ontology (GO) analysis

For the generation of curated ubiquitome, proteins identified in this study were integrated with site-level ubiquitination data from ten previously published datasets (Supplemental Table 3). GO analysis was performed using the ShinyGO software (Ge et al., 2020).

## Acknowledgments

We would like to thank Robert Chalkley and Peter Baker for providing technical expertise. We would also like to thank Prof. Neil Olszewski for sharing the *spy*-4 mutants. We thank Andrea Mair and Prof. Dominique Bergmann for sharing the TurboID constructs.

## Author contributions

S.-L.X. conceptualization; S.K., T.G, Biotinylated protein and peptide-level experiments; S.K., A.V.R., TMT experiments; A.V.R., R.S., WT/*spy-4*/*sec-5* SILIA experiments. S.K., ubiquitination experiments. A.V.R., S.K,. T.G., R.S., data analysis; S.K., D.B, constructs and genetic analysis. W.N. IP-MS analysis. S.K., A.V.R., T.G., Figures. S.K., T.G., S.-L.X. manuscript writing.

## Supporting materials

**Supplementary Fig. S1.**
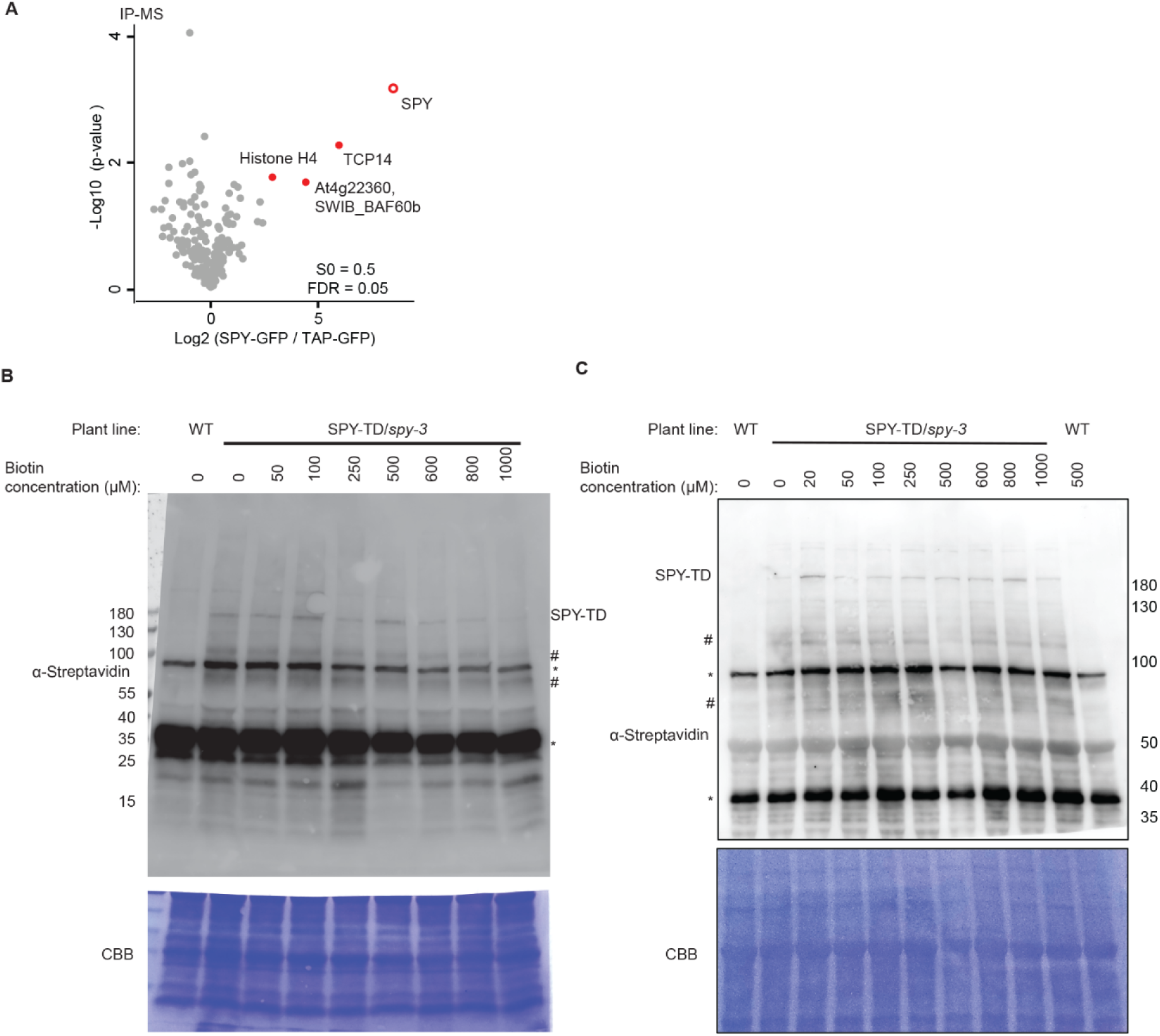
SPY-GFP IP-MS and streptavidin western blot analysis of SPY-TD. (**A**) Volcano plot of SPY-GFP IP-MS. Immunoprecipitation followed by mass spectrometry identified only a small set of specifically enriched proteins, including the bait SPY, the known target TCP14, AT4G22360 (a SWIB complex BAF60b domain–containing protein), and one newly detected interactor, histone H4. (**B**–**C**) Streptavidin western blots reveal very few biotinylated bands and only modest increases in biotinylation with increasing biotin concentration. Ten-day-old (B) and five-day-old (C) SPY-TD seedlings were treated with 0–1000 μM biotin for 1 h. WT seedlings treated with 0 or 500 μM biotin for 1 h served as controls. Total protein extracts were separated on 4–20% (B) or 8% (**C**) SDS–PAGE gels. Streptavidin-HRP immunoblots are shown (top), with Coomassie Brilliant Blue (CBB) staining as a loading control (bottom). Asterisks (*) indicate endogenous biotinylated proteins. The SPY-TD band and two highlighted biotinylated bands are marked with (#).

**Supplementary Fig. S2.**
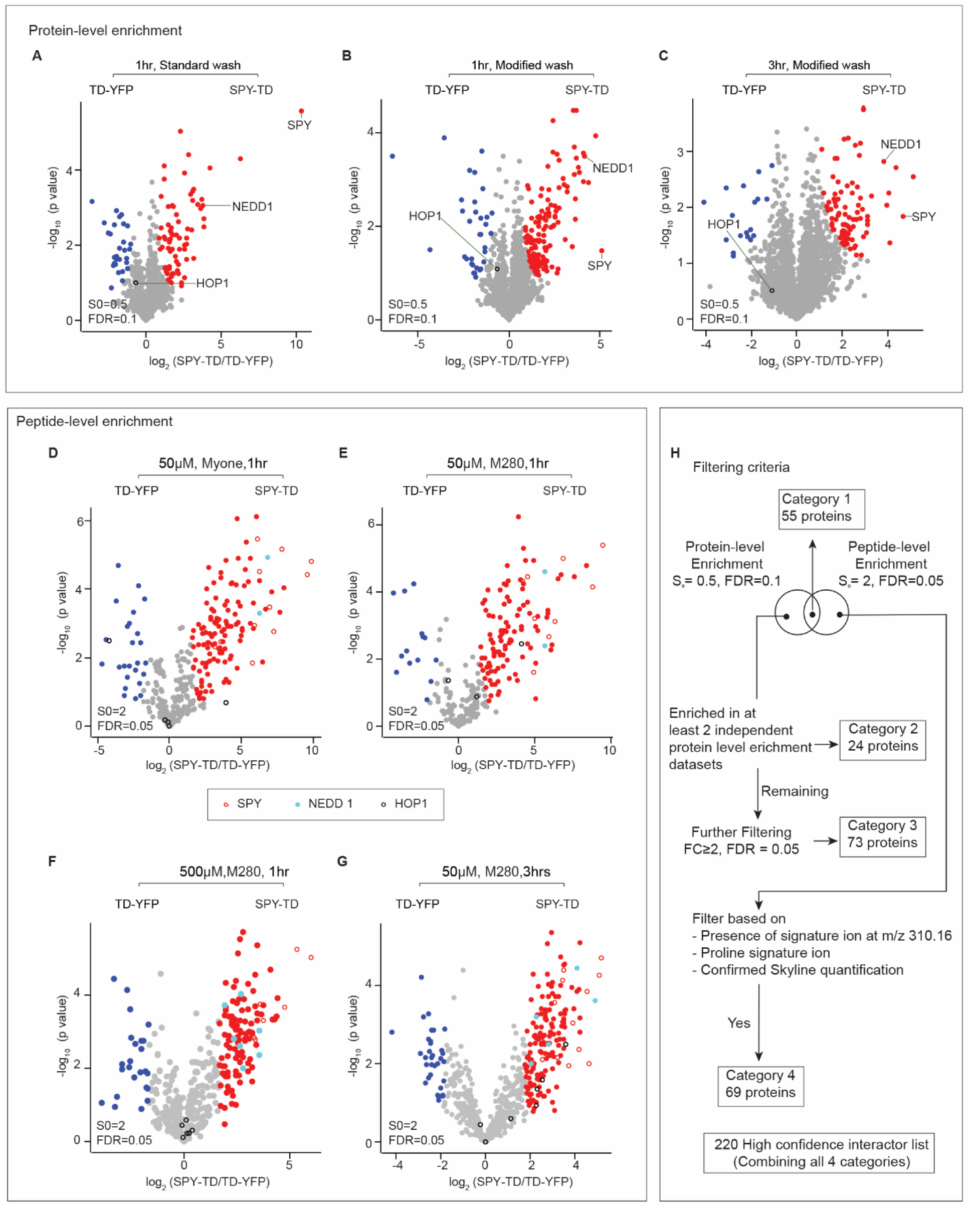
Systematic optimization of SPY-TD labeling conditions and bioinformatics pipeline for high-confidence interactor identification. (**A**–**C**) Optimization of protein-level enrichment. Volcano plots showing protein enrichment across various experimental parameters, including standard vs. modified bead-washing protocols (A vs. B) and varying biotinylation durations (B vs. C). Proteins that were either non-enriched or preferentially enriched in the TD-YFP control are indicated by grey and blue circles, respectively. While the 3-hour biotin treatment increases overall protein biotinylation in both the bait and control samples, certain SPY targets are masked by the increased background in the control. (**D**–**G**) Optimization of peptide-level enrichment. Volcano plots comparing the performance of different streptavidin bead types (MyOne (D) vs. M280 (E)), biotin concentrations (E vs. G), and biotinylation durations (F vs. G). Key proteins are highlighted: SPY (bait), NEDD1 (positive control), and HOP1 (a known nuclear protein susceptible to TD-mediated biotinylation). (**H**) Filtering pipelines to generate high confidence SPY-TD interactor list.

**Supplementary Fig. S3.**
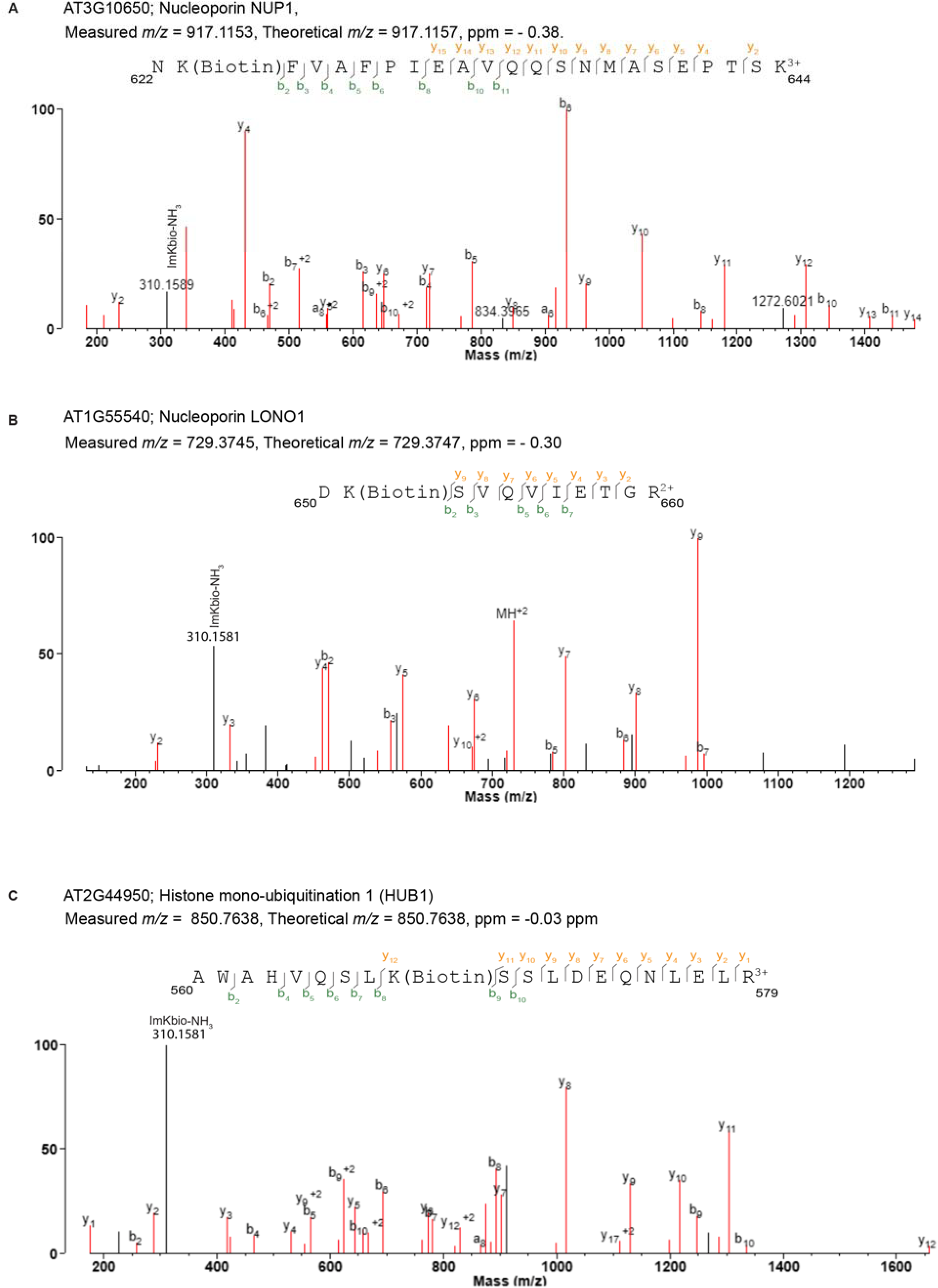
Representative MS/MS spectra and site-specific identification of biotinylated peptides in NUP1, LONO1, and HUB1. (**A**–**C**) MS/MS validation of biotinylation sites. Representative fragmentation spectra unambiguously confirming the amino acid sequences of peptides derived from NUP1, LONO1, and HUB1. The signature immonium-K-biotin–NH₃ ion (*m/z* 310.16; C₁₅H₁₉N₂O₂S), an immonium derivative of biotinylated lysine, is highlighted in each spectrum as a diagnostic marker for the biotin modification. Precise site assignment is confirmed by the presence of site-determining fragment ions (e.g., b- and y-series) that exhibit the characteristic mass shift associated with biotinylation.

**Supplementary Fig. S4.**
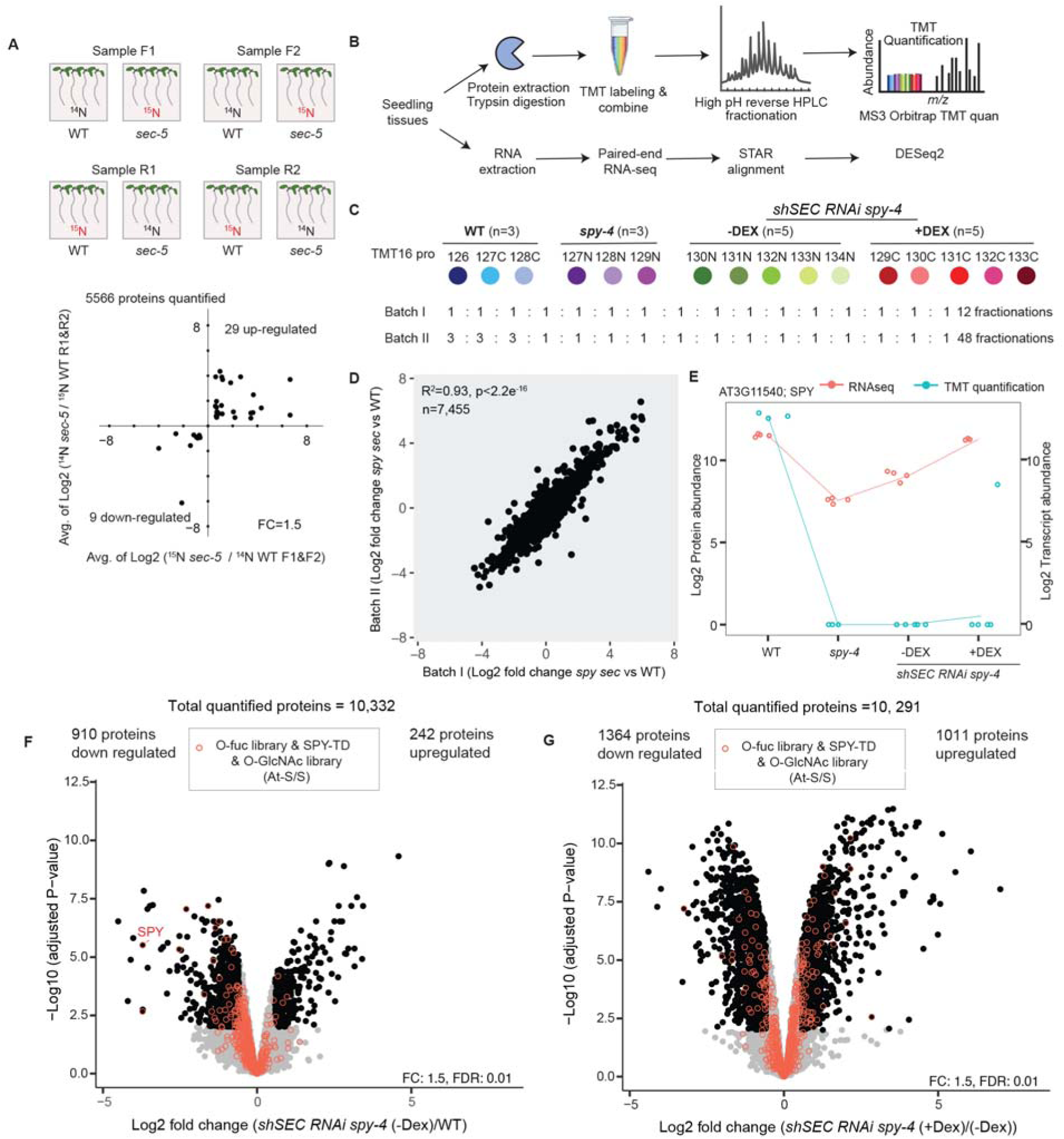
Comparative SILIA analysis of sec-5 and integrated TMT/RNA-seq quantification workflow. (**A**) SILIA-based quantification of WT and *sec-5* mutants. Scatter plot showing protein-level differences between WT and *sec-5*. Consistent with the subtle mutant phenotype, only 38 proteins met the differential threshold (fold change [FC] ≥1.5); non-significant proteins are omitted for clarity. The *x*-axis represents the average forward-labeling ratio, and the *y*-axis represents the average reverse-labeling log_2_ ratio. (**B**–**C**) Schematic overview of the TMT labeling experiment and RNA-seq analyses. Panel (B) summarizes the experimental workflow, and panel (C) details the TMT channels, mixing proportions, and fractionation schemes for batch I and batch II. (**D**) Inter-batch consistency. Comparison of batch I and batch II TMT quantifications shows strong reproducibility between these two approaches. (**E**) SPY protein and RNA abundance across genotypes. Data are presented as log2 protein abundance and log2 normalized RNA counts in WT, *spy-4*, and *shSEC RNAi spy-4* (± DEX). Notably, SPY RNA levels are elevated in the mock-treated *shSEC RNAi spy-4* line and further upregulated upon DEX treatment, suggesting that a reduction in SEC levels triggers a compensatory upregulation of SPY expression (*spy-4* T-DNA insertion is in the 5′ UTR). At the proteomic level, only a single peptide was identified for the SPY protein; SPY remains undetectable in *spy-4* and both mock- and DEX-treated *shSEC RNAi spy-4* samples. One TMT channel for this specific peptide is indicated as an outlier due to suspected contamination. (**F**–**G**) Differential protein abundance in the inducible *shSEC RNAi spy-4* line. Volcano plots showing protein-level changes in *shSEC* RNAi *spy-4* mock treatment compared to WT (F), and following DEX treatment compared to mock treatment (G).

**Supplementary Fig. S5.**
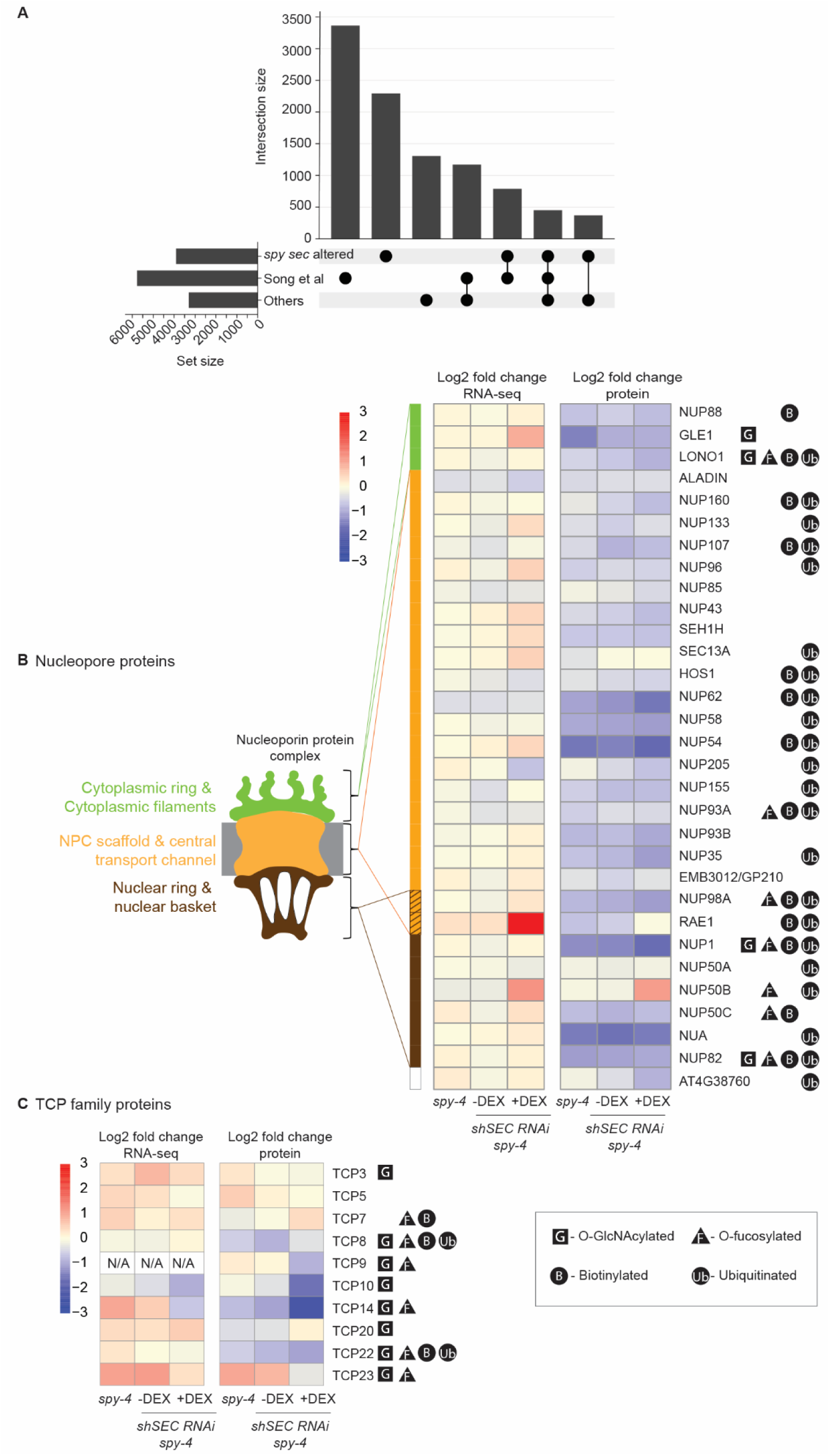
Potential crosstalk between O-glycosylation and ubiquitination among down-regulated proteins in *spy sec* mutants. (**A**) UpSet plot illustrating the overlap between proteins with reduced abundance in the *spy sec* double mutant and two comprehensive ubiquitination datasets. These include: (1) the large-scale ubiquitome identified by Song et al. (2025), and (2) a secondary combined dataset comprising nine previously published studies along with our in-house plant ubiquitome data. (**B**–**C**) O-glycosylation and ubiquitination status of nucleoporins (B) and TCP family proteins (C) in *spy-4* and *shSEC RNAi spy-4* mutants (± DEX). Respective labels indicate O-GlcNAcylation, O-fucosylation, biotinylation, and ubiquitination status. Transcriptomic and proteomic data are presented side-by-side.

**Supplementary Fig. S6.**
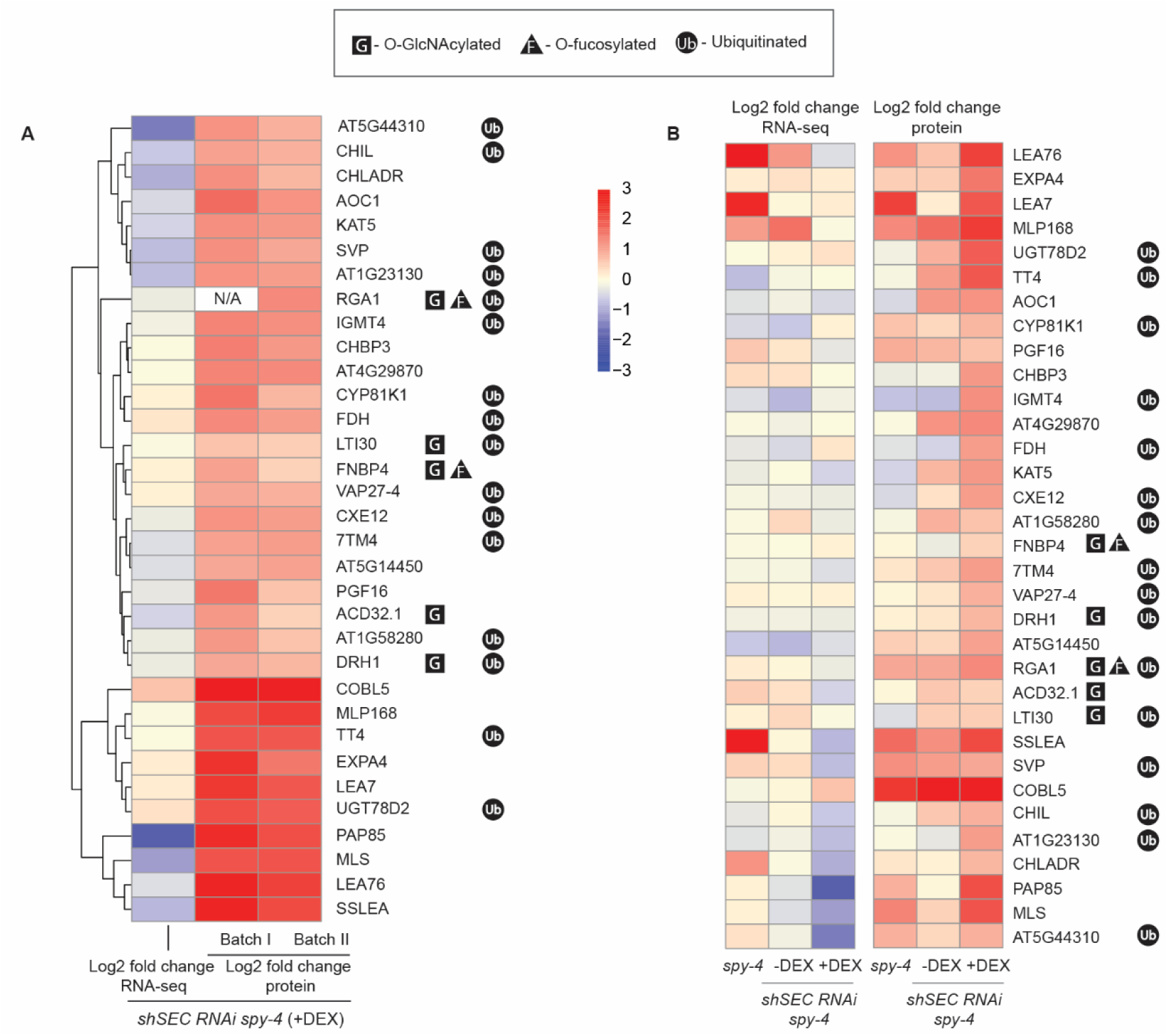
Potential crosstalk between O-glycosylation and ubiquitination among up-regulated proteins in *spy sec* mutants. (**A**) Heatmap illustrating proteins with increased abundance (stabilized) in the *spy sec* mutant relative to WT across two TMT proteomics datasets (Batch I and Batch II). (**B**) Heatmaps comparing RNA and protein abundance for the candidates identified within *spy-4* and *SEC RNAi spy-4* lines under mock and DEX-treated conditions. Respective labels indicate the O-GlcNAcylation, O-fucosylation, and ubiquitination status of each protein. Transcriptomic and proteomic data are presented side-by-side.

**Supplemental Table S1.** Dataset of SPY-GFP IP-MS and SPY-TD PL-MS.

**Supplemental Table S2.** Quantitative proteomic analysis (SILIA and TMT) comparing WT, *spy-4, sec-5, shSEC RNAi spy-4* mutants (± DEX).

**Supplemental Table S3.** Compiled analysis of the ubiquitome.

## Data availability

The mass spectrometry proteomics data have been deposited to the ProteomeXchange Consortium via the PRIDE partner repository, with accession number: PXD050677, PXD050678, PXD050679. All other data are available from the corresponding author on reasonable request. The spectra for the biotinylated peptide identification can be viewed with search key sxu814aioe using MS-viewer (Baker and Chalkley, 2014).

